# Candidate Phyla Radiation Roizmanbacteria from hot springs have novel, unexpectedly abundant, and potentially alternatively functioning CRISPR-Cas systems

**DOI:** 10.1101/448639

**Authors:** Lin-Xing Chen, Basem Al-Shayeb, Raphael Meheust, Wen-Jun Li, Jennifer A. Doudna, Jillian F. Banfield

## Abstract

The Candidate Phyla Radiation (CPR) comprises a huge group of bacteria that have small genomes that rarely encode CRISPR-Cas systems for phage defense. Consequently, questions remain about their mechanisms of phage resistance and the nature of phage that infect them. The compact CRISPR-CasY system (Cas12d) with potential value in genome editing was first discovered in these organisms. Relatively few CasY sequences have been reported to date, and little is known about the function and activity of these systems in the natural environment. Here, we conducted a genome-resolved metagenomic investigation of hot spring microbiomes and recovered CRISPR systems mostly from Roizmanbacteria that involve CasY proteins that are divergent from published sequences. Within population diversity in the spacer set indicates current *in situ* diversification of most of the loci. In addition to CasY, some Roizmanbacteria genomes also encode large type I-B and/or III-A systems that, based on spacer targeting, are used in phage defense. CRISPR targeting identified three phage represented by complete genomes and a prophage, which are the first reported for bacteria of the Microgenomates superphylum. Interestingly, one phage encodes a Cas4-like protein, a scenario that has been suggested to drive acquisition of self-targeting spacers. Consistent with this, the Roizmanbacteria population that it infects has a CRISPR locus that includes self-targeting spacers and a fragmented CasY gene (fCasY). Despite gene fragmentation, the PAM sequence is the same as that of other CasY reported in this study. Fragmentation of CasY may avoid the lethality of self-targeting spacers. However, the spacers may still have some biological role, possibly in genome regulation. The findings expand our understanding of CasY diversity, and more broadly, CRISPR-Cas systems and phage of CPR bacteria.

## Introduction

The Candidate Phyla Radiation (CPR) comprises a huge fraction of Domain Bacteria. The scale of the radiation remains unclear, but it may include as much as 26-50% of all bacterial diversity (Hug et al. 2016; Parks et al. 2017; Schulz et al. 2017). The CPR bacteria uniformly have small genomes (often ~1 Mbp) and limited biosynthetic capacity (Brown et al. 2015; Anantharaman et al. 2016; Hug et al. 2016; Castelle and Banfield 2018). Most are thought to be symbionts, in some cases cell surface attached (episymbionts), that depend on other bacteria for basic cellular building blocks (for review, see (Castelle and Banfield 2018)).

A previous meta-analysis found that only 2.4% of organisms from the Parcubacteria (OD1) and Microgenomates (OP11) superphyla encode CRISPR-Cas systems in their genomes, as compared to 47.4% in archaea and 24.4% in non-CPR bacteria (Burstein et al. 2016). The authors noted that when CRISPR-Cas systems occur in CPR bacteria they tend to be different from those found in other bacteria. Four genomes from Dojkabacteria (WS6), Parcubacteria (OD1) and Roizmanbacteria were previously recognized to encode CRISPR-Cas12a (Cpf1) systems (Zetsche et al. 2015), and more recently, six genomes were reported encoding a newly recognized compact CasY effector enzyme that has genome editing potential (Burstein et al. 2017).

Several potential explanations for the low frequency of CRISPR-Cas systems in CPR bacteria have been suggested (Burstein et al. 2016). Small genome size may favor use of more compact restriction-modification systems for phage defense and low ribosome content may preclude sufficiently fast-acting CRISPR-Cas systems required for effective interference (Burstein et al. 2016). Symbiotic lifestyles, characterized by close association between multiple cells and a host cell, could lead to higher phage densities, which may cause selection of defense systems other than CRISPR-Cas (Westra et al. 2015). It has also been suggested that CPR bacteria may not have the RecBCD mechanism identified in non-CPR Bacteria to curtail self-targeting spacer acquisition (Levy et al. 2015; Castelle et al. 2018).

As few phage that infect CPR bacteria have been reported (Paez-Espino et al. 2016; Dudek et al. 2017), it is difficult to know how common phage that infect these bacteria might be. Phage particles in the process of infecting CPR bacterial cells have been observed via cryogenic electron microscopy (Luef et al. 2015). However, the sequences of phage associated with CPR bacteria are unusually difficult to identify in metagenomic datasets, in part due to the lack of CRISPR spacers that could be used to link them to host cells via CRISPR targeting (Andersson and Banfield 2008). Further, like phage, CPR genomes encode a very high proportion of novel proteins (Castelle and Banfield 2018), which obscures identification of potential prophage regions. Finally, phage structural proteins may be too divergent from those of well-studied phage to be identified. To date, phage have only been reported for bacteria from two CPR phyla, Absconditabacteria (previously SR1) and Saccharibacteria (previously TM7) (Paez-Espino et al. 2016; Dudek et al. 2017). Thus, there is a potentially huge knowledge gap related to the existence and diversity of CPR phage. This motivates the search for new CPR genomes with CRISPR-Cas systems that could potentially provide links to additional examples of phage that replicate in these bacteria.

In the current study, we investigated the microbiomes of a series of hot springs in Tibet. CPR bacteria are relatively abundant in these thermal environments, and some of their genomes encode interesting and unusual CRISPR-Cas systems. Although uncommon overall, CRISPR-Cas systems are surprisingly frequently encoded in the genomes of members of the Roizmanbacteria, and multiple different systems coexist in some genomes. We identified many new examples of systems based on CasY and uncovered an intriguing example of a locus with self-targeting spacers and a fragmented CasY gene. We identified CPR phage for which complete, curated genomes were reconstructed, as well as prophage in other genomes. Thus, our analyses provide new insights into CPR biology, their phage and the diversity of the relatively unstudied CRISPR-CasY system.

## Materials and methods

### Study site, sampling and physicochemical determination

Hot spring (40.8 - 84.9 °C) sediment samples were collected from Tibet Plateau (China) in August 2016 Supplementary Table 1). As described previously (Song et al. 2012), sediment samples were collected from the hot spring pools using a sterile iron spoon into 50 ml sterile tubes, transported to the lab on dry ice, and stored at −80 °C for DNA extraction. Temperature, dissolved oxygen (DO) and pH were determined *in situ* and the other physicochemical parameters were analyzed in the laboratory (Supplementary Table 1).

**Table 1.**
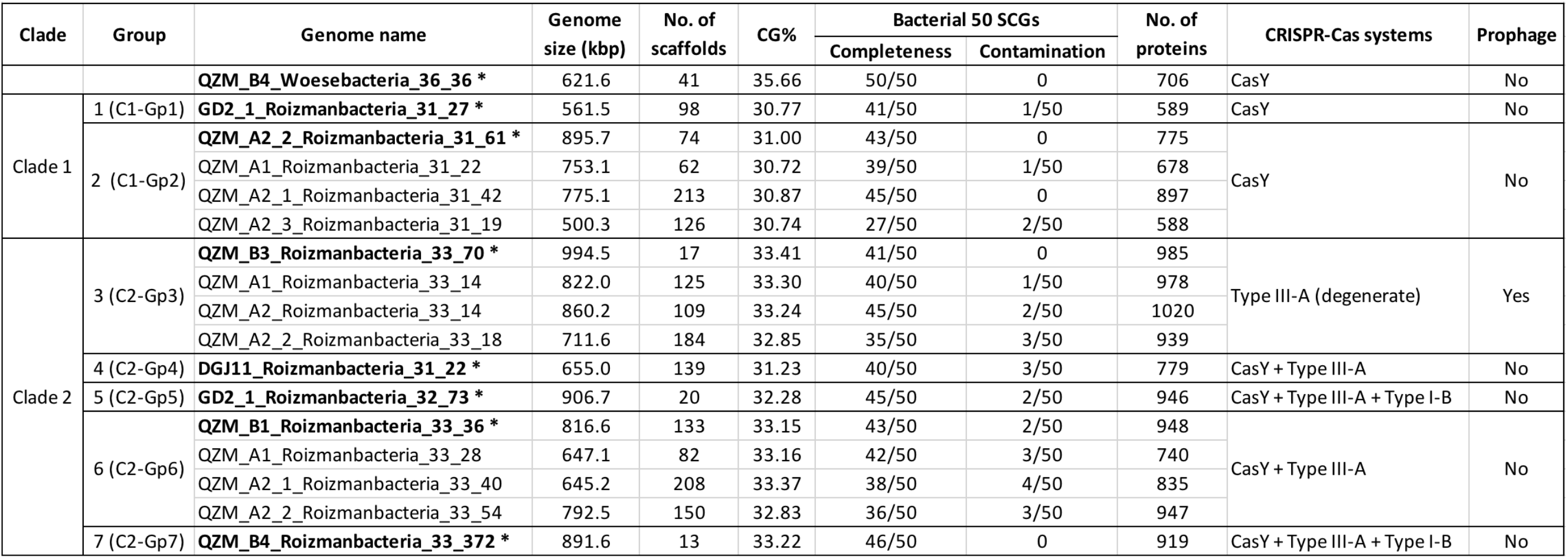
Summary of Roizmanbacteria and Woesebacteria genomes reconstructed in this study. *Representative genome of each group used in phylogenetic analyses. Figure 1 and Supplementary Figure 1 provide phylogeny and clade information.

### DNA extraction, sequencing, quality control and metagenomic assembly

Genomic DNA was extracted from sediment samples using the FastDNA SPIN kit (MP Biomedicals, Irvine, CA) according to the manufacturer’s instructions. The DNA samples were purified for library construction, and sequenced on an Illumina HiSeq2500 platform with PE (paired-end) 150 bp kits. The raw data of each metagenomic dataset were filtered to remove Illumina adapters, PhiX and other Illumina trace contaminants with BBTools, and low quality bases and reads using Sickle (version 1.33; https.github.com/najoshi/sickle). The high-quality reads of each sample were assembled using metaSPADES (version 3.10.1) (Bankevich et al. 2012) with a kmer set of 21, 33, 55, 77, 99, 127.

### HMM-based search of CasY proteins and confirmation of CRISPR-CasY system

The six CasY proteins reported previously (Burstein et al. 2017) were aligned using Muscle (Edgar 2004), and filtered to remove those columns comprising 95% or more gaps with TrimAL (Capella-Gutiérrez, Silla-Martínez, and Gabaldón 2009). A HMM model was built based on the filtered alignment using hmmbuild 2 (Eddy 1998) with default parameters, hmmsearch was used to search all the proteins predicted by Prodigal from scaffolds. Those hits with an e-value < 10-5 were manually checked, and the online tool CRISPRs finder (Grissa et al. 2008) was used to identify the Cas1 protein and CRISPR loci. Only those scaffolds detected with CasY, Cas1 and CRISPR array were retained for further analyses. Other CRISPR-Cas systems identified in these genomes based on the presence of Cas proteins and CRISPR arrays were also analyzed in this study.

### Extension and manual curation of CasY scaffolds

Those scaffolds with partial CasY representatives were manually extended as follows: (1) mapping the high quality reads to the corresponding scaffolds using bowtie2 with default parameters; (2) filtering the mapping files using mapped.py (part of the ra2 suite) to remove those PE reads with two or more mismatches to the assembled scaffold across both reads combined; (3) importing the filtered mapping files into Geneious and mapping using the “Map to Reference” function; (4) extending the scaffolds at the partial CasY protein ends; (5) performing the first 4 steps again (multiple times if necessary) until full length CasY proteins were obtained.

The extended scaffolds and other full-length CasY scaffolds were checked for any potential assembly errors using ra2.py (https://github.com/christophertbrown/fix_assembly_errors/releases/tag/2.00), the general strategy was described previously (Brown et al. 2015). Errors reported as unresolved by ra2.py were fixed manually in Geneious using unplaced paired reads that were mapped to the scaffolding gaps.

### Coverage calculation, genome binning, genome curation and completeness assessment

The high quality reads were mapped to the corresponding assembled scaffolds using bowtie2 with default parameters and the coverage of each scaffold calculated as the total number of bases mapped to it divided by its length. For each sample, scaffolds over 2500 bp were assigned to preliminary draft genome bins using MetaBAT with default parameters, considering both tetranucleotide frequencies (TNF) and scaffold coverage information. The clustering of scaffolds from the bins and the unbinned scaffolds was visualized using ESOM with a min length of 2500 bp and max length of 5000 bp as previously described (Dick et al. 2009). Misplaced scaffolds were removed from bins and unbinned scaffolds whose segments were placed within the bin areas of ESOMs were added to the bins. Scaffolds ≥ 1000 bp from each sample were uploaded to ggKbase (http://ggkbase.berkeley.edu/). The ESOM-curated bins with interesting CasY-bearing scaffolds were further evaluated based on consistency of GC content, coverage and taxonomic information, and scaffolds identified as contaminants were removed. The genome bins with CRISPR-CasY systems were curated individually to fix local assembly errors using ra2.py, as described above. A total of 50 single copy genes (SCGs) that are commonly detected in CPR bacteria (Supplementary Table 2) were used to evaluate genome completeness.

### Gene prediction and metabolic prediction

The protein-coding genes of the curated genomes (see above) were predicted using Prodigal (-m single) (Hyatt et al. 2010), and searched against KEGG, UniRef100 and UniProt for annotation, and metabolic pathways were reconstructed. The 16S rRNA genes were predicted based on HMM models, as previously described (Brown et al. 2015). The ribosome binding site sequence was obtained via the Prodigal gene prediction results.

### CRISPR loci reconstruction and spacer identification

For all the confirmed CRISPR-CasY and other CRISPR-Cas systems, the quality reads were aligned to the scaffolds from the corresponding sample using bowtie2 with default parameters (Brown et al. 2015; Langmead and Salzberg 2012). Any unmapped reads of read pairs were mapped to the scaffolds in Geneious using the function of “Map to Reference”, then the CRISPR loci were manually reconstructed, allowing for spacer set diversification and loss of spacer-repeat units in some cells. Thus, it was possible to place most reads in an order that reflects the locus evolutionary history. For each CRISPR locus, all the reads that mapped were extracted, and spacers between two direct repeats were used for target searches (see below).

### Spacers target search and identification of (pro)phage scaffolds

All the spacer sequences from each CRISPR locus were dereplicated, then the sequences were searched against scaffolds from related samples using BLASTn with the following parameters: -task blastn-short, -dust no, -word_size 8. Those scaffolds with 0 mismatch and 100% alignment coverage to one or more spacers were manually checked for phage-specific proteins, including capsid, phage, virus, prophage, terminase, prohead, tape measure, tail, head, portal, DNA packaging, as described previously (Dudek et al. 2017).

### In silico determination of protospacer adjacent motif (PAM)

To determine the PAM of the CRISPR-CasY systems in Roizmanbacteria genomes, for each CRISPR spacer with a target in two complete phage genomes from QZM (see results), the upstream 5 bp and downstream 5 bp of the targeted DNA strand were searched manually and the PAM was determined and visualized using Weblogo (Crooks et al. 2004). The PAM analyses for other CRISPR-Cas systems analyzed in this study were performed in the same way.

### Phylogenetic analyses

Phylogenetic analyses were performed using (1) 16 ribosomal proteins (16 RPs) and (2) 16S rRNA genes of genomes of interest with CRISPR-CasY and/or other CRISPR systems (Table 1), (3) CasY proteins, (4) Cpf1 proteins, and (5) capsid proteins of CPR (pro)phage:

1. 16 RPs analyses: After preliminary classification based on the ribosomal protein S3 (rpS3) taxonomy, reference genomes were downloaded from NCBI (131 in total) and dereplicated using dRep (“-sa 0.95 -nc 0.5”) (Olm et al. 2017). A higher similarity threshold was used to perform dereplication of newly reconstructed genomes from hot spring sediment samples (“-sa 0.99 -nc 0.5”), to clarify the overall diversity. The 16 RPs (i.e., L2, L3, L4, L5, L6, L14, L15, L16, L18, L22, L24, S3, S8, S10, S17 and S19) were predicted from all the dereplicated genomes.
2. 16S rRNA genes sequences: The 16S rRNA genes were predicted from all the dereplicated genomes (see above) using HMM-based searches (Brown et al. 2015). All the insertion sequences with lengths > 10 bp were removed.
3. CasY proteins: all partial and full length CasY proteins from confirmed CRISPR-CasY systems in this study and the previously reported CasY proteins were included in a phylogenetic tree, with c2c3 proteins as the outgroup.
4. Cas12a (Cpf1) proteins: the Cas12a proteins in NCBI and our dataset were identified and used to construct a tree with Cas12c (C2c3) proteins as the outgroup.
5. CPR (pro)phages: the capsid protein was used as a marker to build phylogenetic trees for CPR (pro)phage. The capsid proteins identified in this study were searched against the NCBI RefSeq Phage Capsid proteins, the first 5 blast hits were used as reference proteins, along with those in previously reported in CPR phage genomes (Paez-Espino et al. 2016; Dudek et al. 2017).

For tree construction, protein sequences datasets were aligned using Muscle (Edgar 2004). The 16S rRNA gene sequences were aligned using the SINA alignment algorithm (Edgar 2004; Pruesse, Peplies, and Glöckner 2012) through the SILVA web interface (Pruesse et al. 2007). All the alignments were filtered using TrimAL (Capella-Gutiérrez, Silla-Martínez, and Gabaldón 2009) to remove those columns comprising more than 95% gaps. For the 16 RP, ambiguously aligned C and N termini were removed and the amino acid sequences, which were concatenated in the order as stated above (alignment length, 2654 aa). The phylogenetic trees were reconstructed using RAxML version 8.0.26 with the following options: -m PROTGAMMALG (GTRGAMMAI for 16S rRNA phylogeny) -c 4 -e 0.001 -# 100 -f a (Capella-Gutiérrez, Silla-Martínez, and Gabaldón 2009; Stamatakis 2014). All the trees were uploaded to iTOL v3 for visualization and formatting (Letunic and Bork 2006).

### Data availability

The reconstructed CPR and their infecting phage genomes reported in the current study were deposited at NCBI within BioProject PRJNA493250 (BioSample SUB4567433), under the accession numbers of xxx-xxx. The unbinned scaffolds with CRISPR-CasY system were deposited under the NCBI accession numbers of xxx-xxx. All genomic data can be explored and downloaded from ggKbase (https://ggkbase.berkeley.edu/Tibet_CRISPR_CasY/organisms) following publication of this manuscript. Note that registration by provision of an email address is required prior to data download.

## Results

### Newly reconstructed Roizmanbacteria and Woesebacteria genomes with CRISPR-Cas systems

CPR bacteria collectively accounted for up to 43.1% of the analyzed hot spring communities (Supplementary Figure 1. We selected 17 genomes that encode CRISPR-Cas systems for curation (Figure 1a, Table 1). Based on rpS3 protein taxonomic analysis, one Woesebacteria genome and 12 Roizmanbacteria genomes encode CRISPR-CasY systems, and four other Roizmanbacteria genomes encode only type III-A CRISPR-Cas systems. Both of these phylum-level groups place within the Microgenomates (OP11) (Hug et al. 2016; Brown et al. 2015). Phylogenetic analyses based on 16 RPs with published Roizmanbacteria genomes (43 dereplicated in total) indicated the divergence of the newly reconstructed Roizmanbacteria from previously published genomes (Figure 1a). The new Roizmanbacteria genomes were assigned to two distinct classes based on their 16S rRNA gene sequences (Yarza et al. 2014) and/or average nucleotide identity (ANI) (Figure 1a, Supplementary Figure 2). Five of the genomes represent two different strains, with an ANI of 98.39% (clade 1; Figure 1a), and the other 11 genomes belong to the same family (clade 2; Figure 1a). Genomes in clade 1 and 2 were assigned to groups (Figure 1a, Table 1).

**Figure 1.**
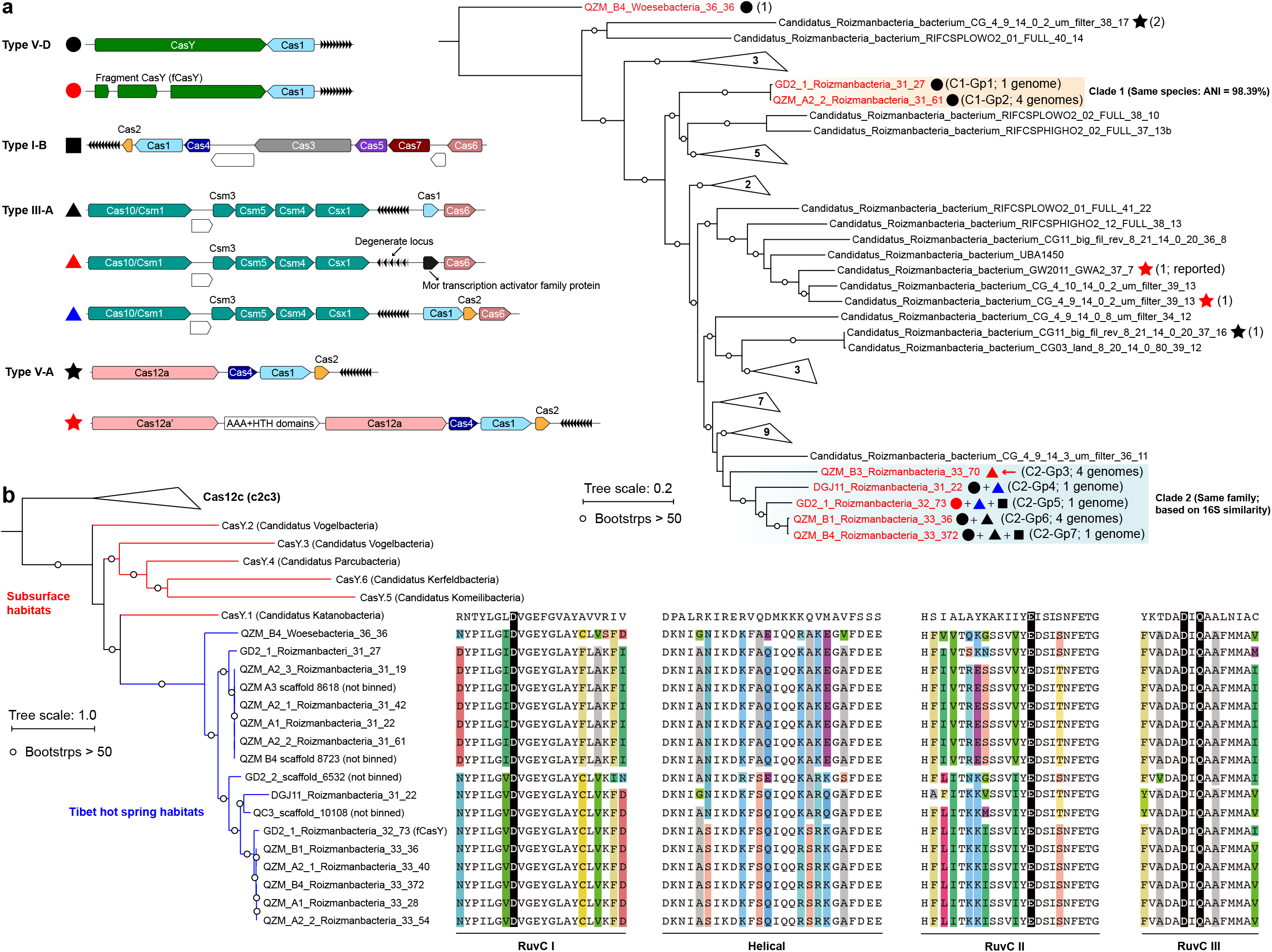
**Roizmanbacteria and Woesebacteria genomes encode CRISPR-CasY and/or other CRISPR-Cas systems**. (a) CRISPR systems detected in genomes of CPR bacteria (top left), the phylogenetic classification of which was established based on concatenated sequences of 16 ribosomal proteins (top right). Included in the analyses are the dereplicated representatives of Roizmanbacteria and Woesebacteria genomes from this study (in red) and Roizmanbacteria genomes from NCBI (in black). CRISPR system types in each genome are indicated by the symbols after the genome names, and the number of non-redundant genomes is shown in brackets. Those clades without CRISPR-Cas are collapsed, and the number of genomes are shown (see Supplementary Figure 1 for the uncollapsed tree). The red arrow indicates the presence of a restriction-modification system, the hypothetical proteins are shown in white. (b) Phylogenetic analyses of CasY proteins, including those previously reported and those identified in this study. The local alignment of conserved motifs of CasY protein, including RuvC-I, -II, -III and helical, are shown. The catalytic residues are shown by white letters on a black background; for other residues, backgrounds of different colors are used if the amino acids are inconsistent among those CasY identified in this study.

**Figure 2.**
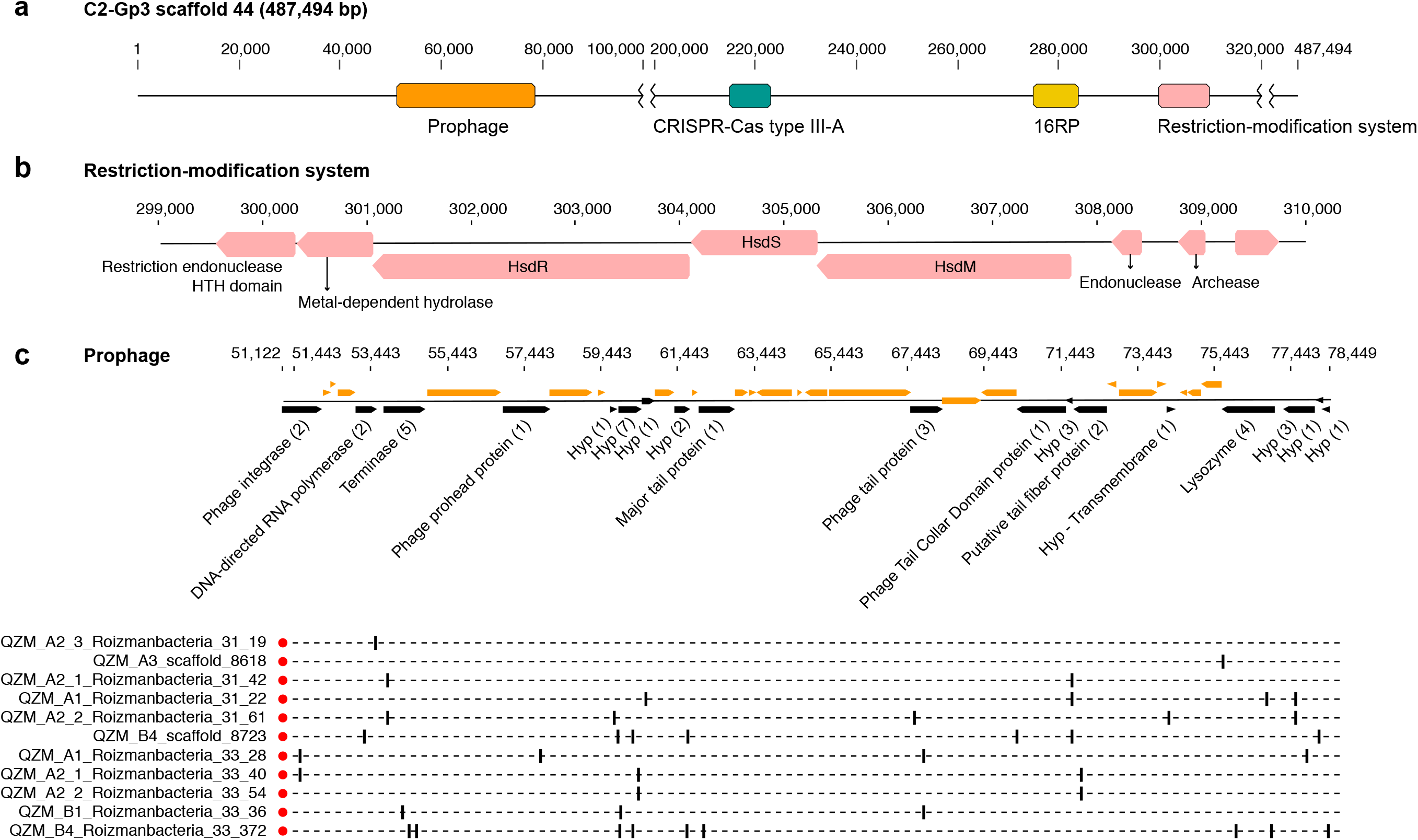
**Prophage and restriction modification systems are detected in the genomes of Roizmanbacteria C2-Gp3**. (a) Scaffold 44 includes prophage, a restriction-modification system and an apparently degenerate Type III-A CRISPR-Cas system. (b) The proteins in the restriction-modification system shown in (a). (c) Prophage genes targeted by spacers are shown in black and the number of spacers targeting each open reading frame is listed in brackets following the annotation. Genes not targeted by CRISPR spacers are shown in orange (top panel). The genome affiliations of spacers targeting the prophage are indicated in the bottom panel.

### CRISPR-CasY detected in Roizmanbacteria and Woesebacteria genomes

We identified 69 CasY candidates (see methods), 17 of which are on scaffolds with a Cas1 protein and CRISPR locus (Supplementary Table 3). Of these, 12 scaffolds could be assigned to Roizmanbacteria genomes and one to a Woesebacteria genome (Table 1). The other four scaffolds with CRISPR-CasY systems could not be binned, but were also included in our analyses (Figure 1b).

The CRISPR-CasY systems from of Roizmanbacteria and Woesebacteria have a different architecture than those reported previously (Burstein et al., 2017), with CasY and Cas1 proteins on the same side of the CRISPR locus (Figure 1a). The Roizmanbacteria CasY proteins have similar lengths of 1252-1256 aa, whereas that found in the Woesebacteria is 1304 aa (Supplementary Table 3), comparable to lengths of previously reported CasY (1153-1287 aa; (Burstein et al. 2017)). Phylogenetic analyses of CasY proteins showed that the newly reported Roizmanbacteria and Woesebacteria sequences are most closely related to CasY.1 from Candidatus Katanobacteria (WWE3; (Burstein et al. 2017)) (Figure 1b).

CasY is an effector protein of Type V CRISPR-Cas systems. To date, all reported Type V CRISPR-Cas systems have RuvC-like nuclease domains (Burstein et al. 2017; Chen and Doudna 2017). Comparative analyses of all CasY proteins reported in this study and CasY.1 with Cpf1, C2c1 and C2c3 references (Shmakov et al., 2015), identified all the catalytic residues within the three conserved motifs of RuvC-I, RuvC-II and RuvC-III (Figure 1b), suggesting the RuvC domains in the new CasY proteins are active nucleases. On the other hand, we detected divergence in other regions of the CasY proteins from different sampling sites (Figure 1b).

### Other CRISPR-Cas systems identified in Roizmanbacteria genomes

A Type III-A system was detected in all 11 clade 2 Roizmanbacteria genomes, seven of which encode more than one type of system (Figure 1a, Supplementary Figure 2). The genomes differ in terms of the presence or absence of Cas1 and Cas2 proteins (Supplementary Table 3), which are used for acquisition of new spacers (Shmakov et al. 2015; Hille et al. 2018; Nuñez et al. 2014). In detail, III-A systems in C2-Gp4 and C2-Gp5 have both Cas1 and Cas2. C2-Gp6 and C2-Gp7 possess Cas1 but not Cas2. Four genomes in C2-Gp3 lack both Cas1 and Cas2 but have a Mor transcription activator family protein (Figure 1a and Supplementary Table 3). However, the CRISPR-Cas system in C2-Gp3 may be non-functional because the repeats are imperfect. A fragment of the C2-Gp3 genomes encodes the 16 ribosomal proteins used for phylogenetic analyses and a restriction-modification system that may instead be used for phage defense (Figures 2a and b).

A Type I-B system was identified in two Roizmanbacteria genomes belonging to the same genus (C2-Gp5 and C2-Gp7), but not in the C2-Gp6 genomes, despite the fact that C2-Gp5 and C2-Gp7 are very closely related to C2-Gp6 (ANI = 99% and 16S similarity = 98.9%). Comparative genomic analyses showed that the Type I-B system is located between genes encoding a secreted cysteine-rich protein and a lamin tail domain protein that are present in both genomes (Supplementary Figure 3b). Two very short hypothetical proteins were detected between the cysteine-rich and lamin tail domain proteins in the C2-Gp6 genomes (Supplementary Figure 3b). However, NCBI BLAST and HMM searches indicate no homology of the hypothetical proteins to any known proteins or functional domains, respectively, and no significant similarity to the Cas proteins of Type I-B systems in the C2-Gp5 and C2-Gp7 genomes.

**Figure 3.**
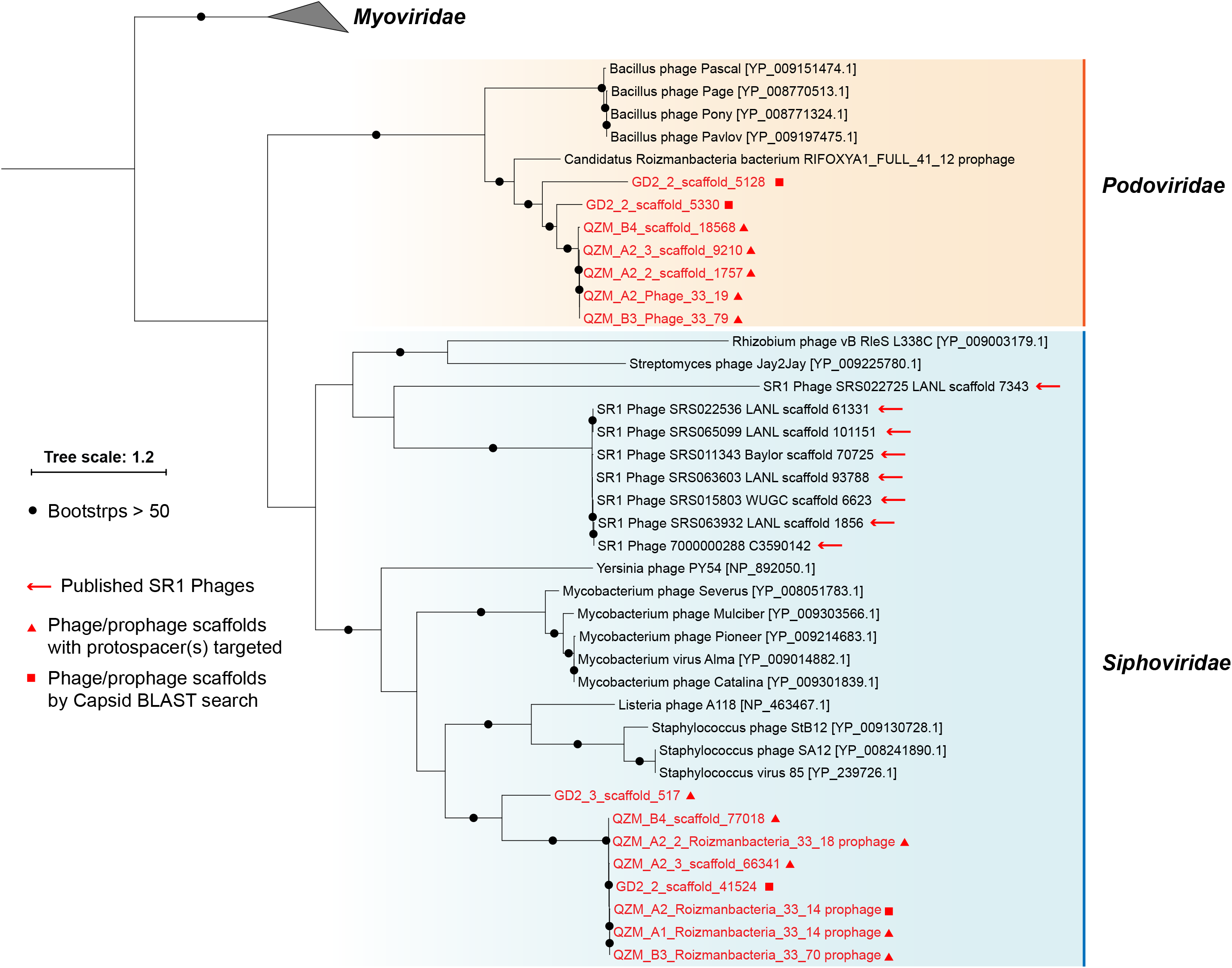
**Phylogeny of capsid proteins used for taxonomic assignment of Roizmanbacteria-infecting phage (or prophage) in this study**. The Roizmanbacteria-infecting phage and prophage are shown in red, and those with spacer targets are indicated by triangles. Squares indicate phage determined to be similar based on their capsid protein sequences. The previously reported Absconditabacteria (SR1) phage are included for comparison.

### CRISPR-Cas12a systems in published Roizmanbacteria genomes

We investigated 131 published Roizmanbacteria genomes available from NCBI to identify all CRISPR-Cas systems that occur in these bacteria (Supplementary Table 4). The CRISPR-Cas12a system (Cpf1), which was identified in one Roizmanbacteria genome (Zetsche et al. 2015), occurred in four Roizmanbacteria genomes from two classes (Figure 1a, Supplementary Figures 2 and 4), one of them in the class containing Roizmanbacteria clade II with type I-B and III-A systems (see above).

**Figure 4.**
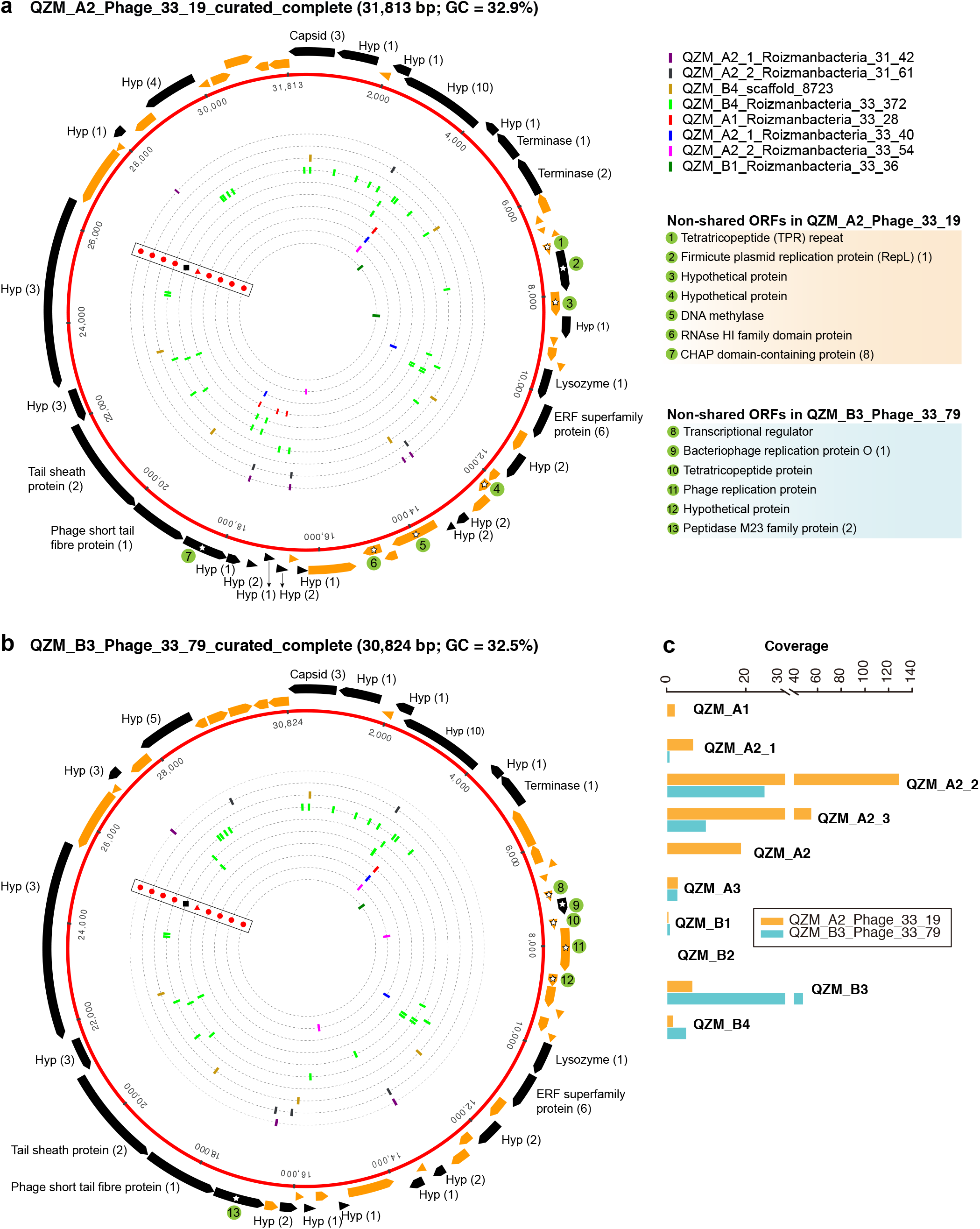
**Complete genomes of Roizmanbacteria-infecting phage**. The red rings represent (a) QZM_A2_Phage_33_19 and (b) QZM_B3_Phage_33_79 phage genomes. The open reading frames (ORFs) are shown outside the genomes, those targeted by at least one spacer are in black (genes not targeted are in orange). The total number of spacers that target each gene is listed in parentheses following the protein annotation. The spacers targeting the phage genome from a given CRISPR-Cas system are indicated by bars on the dotted inner rings (see Figure 1 for CRISPR-Cas system type). Bars are colored by genome of origin (see top right). The non-shared proteins between these two phage genomes are indicated by green circles and numbered, their annotations are shown at the right. Hyp, hypothetical protein. (c) The coverage information of these two phage genomes in QZM-related samples.

Interestingly, the CRISPR-Cas12a systems reported in (Zetsche et al. 2015) from Candidatus Roizmanbacteria bacterium CG_4_9_14_0_2_um_filter_39_13 included two Cas12a proteins. We refer to the one near the CRISPR locus as Cas12a, and the other as Cas12a’ (Figure 1a). Phylogenetic analyses of Cas12a and Cas12a’ proteins (previously reported and identified in this study) indicated those in CPR genomes could be assigned into at least three groups (Supplementary Figure 4a). Group 1 includes the Cas12a proteins from the two genomes with both Cas12a and Cas12a‘, and is highly divergent from other Cas12a proteins. Group 2 includes the Cas12a’ of Candidatus Roizmanbacteria bacterium CG_4_9_14_0_2_um_filter_39_13, along with the Cas12a proteins from another two genomes. Group 3 includes Cas12a’ of Candidatus Roizmanbacteria bacterium GW2011_GWA2_37_7 (Zetsche et al. 2015) and clusters together with Cas12a from non-CPR Bacteria and Archaea.

The RuvC domains (-I, -II, -III) of the CPR Cas12 and Cas12’ group 2 and 3 proteins include all the conserved catalytic residues in (Supplementary Figure 4b). However, in group 1 proteins, the conserved RuvC-II glutamic acid catalytic residue “E” was substituted by asparagine “N”, and in RuvC-III asparagine “N” was substituted to valine “V”. These substitutions suggest that the Cas12a in the systems with both Cas12a and Cas12a’ may not perform cleavage as documented previously (Zetsche et al. 2015).

### Roizmanbacteria-infecting phages from Podoviridae and Siphoviridae

A total of 1118 spacers perfectly targeted (100% match and 100% alignment coverage; see methods) 565 unique scaffolds. Of these, 156 of them were targeted by two or more spacers (153 from the QZM samples of the current study) (Supplementary Table 5). Eleven of the CRISPR spacer-targeted scaffolds encode a phage capsid protein, which was used as a marker for phylogenetic analyses (Figure 3). Five additional scaffolds encoding a similar capsid protein were identified by a BLAST search. Capsid proteins were also predicted from the Absconditabacteria (SR1) phage (8 out of 17 with capsid genes identified) and included in the phylogenetic analyses. The (pro)phage identified in this study as well as the Absconditabacteria phage were assigned to either the *Podoviridae* clade or the *Siphoviridae* clade (Figure 3). The complete Saccharibacteria phage that lacks an identifiable capsid protein (Dudek et al. 2017) is most closely related to *Siphoviridae* phages based on comparison of its terminase with annotated sequences in the NCBI database.

One scaffold (QZM_B3_scaffold_44) from a C2-Gp3 Roizmanbacteria was targeted by multiple spacers. Detailed analyses indicate that this region is a prophage, with a length of approximately 27 kbp (Figure 2c), and is among the first prophage reported in CPR bacterial genomes. This prophage is predicted to encode 40 protein coding genes, including a phage integrase, terminase, prohead protein, major tail protein, tail tape measure protein, tail fiber protein and lysozyme. Nineteen of the ORFs were targeted by 41 CRISPR spacers from CasY-based systems, all of which were from Roizmanbacteria (Figure 1, Figure 4b). BLAST comparison detected highly similar scaffolds in the other three genomes of the C2-Gp3 group (Table 1, Figure 2) and also unbinned scaffolds in QZM_A2_1, QZM_A2_3 and QZM_A3, suggesting that this is a common Roizmanbacteria prophage. However, when reads of other QZM-related samples were mapped to QZM_B3_scaffold_44, the prophage region showed much higher coverage in QZM_B1 and QZM_B4 than the flanking region (Supplementary Figure 5). Further, a subset of reads that circularize the phage genome were detected. These observations indicate that the prophage existed as phage particles in these two samples.

**Figure 5.**
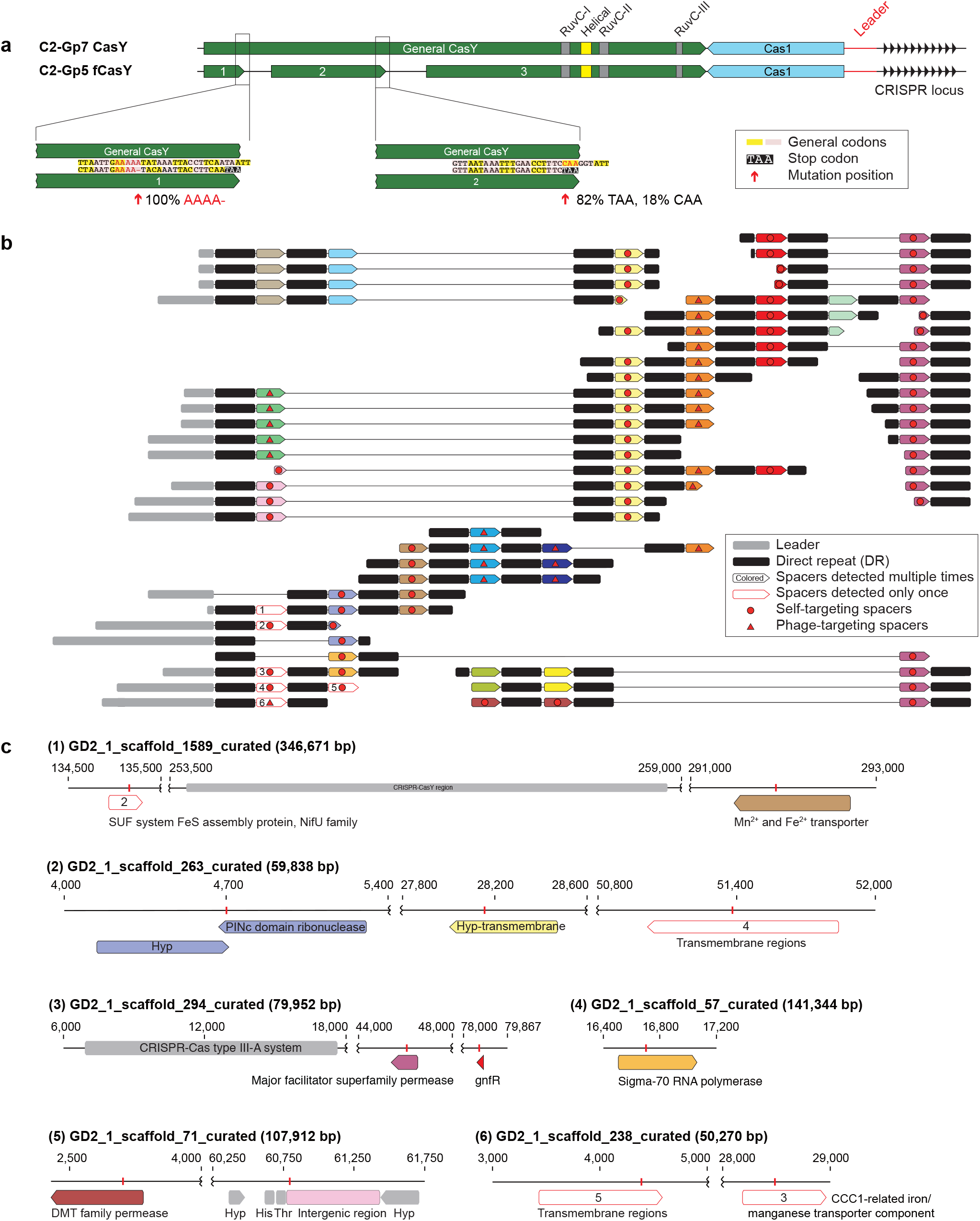
**One Roizmanbacteria genome encodes and unusual CRISPR system with a fragmented CasY (fCasY) protein and self-targeting spacers**. (a) Mutations leading to fragmentation of CasY proteins into three pieces (red arrows) and their incidence in the population, and other features of the locus. (b) The reconstructed CRISPR locus showing the history of spacer acquisition and the distribution of self-targeted spacers (marked by red circles). (c) Scaffolds encoding genes and an intergenic region matching the self-targeting spacers. The targeted genes have the same color as the corresponding spacers in (b), genes targeted by single copy spacers (white in (b)) are indicated by numbers, and CRISPR-Cas systems, tRNA and other genes on the scaffolds are shown in gray.

One putative phage scaffold (Supplementary Table 5) could be circularized, and circularization of the genome was confirmed by paired-end read mapping. The length of complete phage genome QZM_A2_Phage_33_19 is 31,813 bp, with a GC content of 32.9% (Figure 4a). Another two scaffolds (Supplementary Table 5) were manually curated to generate another complete phage genome QZM_B3_Phage_33_79, with a length of 30,824 bp and GC content of 32.5% (Figure 4b). Phage QZM_A2_Phage_33_19 and QZM_B3_Phage_33_79 share high sequence similarity, and they are probably closely related strains.

A total of 53 and 52 open reading frames (ORFs) were predicted from QZM_A2_Phage_33_19 and QZM_B3_Phage_33_79, respectively (Figures 4a and b, Supplementary Figure 6). Of these, 46 shared an average amino acid identity of 98%. ORFs common to both phage encode capsid, terminase, lysozyme and tail proteins. Although these two genomes are highly similar, 7 and 6 non-shared ORFs were detected in QZM_A2_Phage_33_19 and QZM_B3_Phage_33_79, respectively. Among those 13 non-shared ORFs, 4 are related to phage replication (Figures 3a and b), including one replication protein in QZM_A2_ Phage_33_19, and one transcriptional regulator and two replication proteins in QZM_B3_Phage_33_79. We used the divergent region between the two genomes to calculate the coverage of the phage in all QZM-related samples and found that they co-occur in most samples (Figure 4c).

A total of 63 spacers targeted 26 ORFs in QZM_A2_Phage_33_19, and 52 spacers targeted 22 ORFs in QZM_B3_Phage_33_79 (Figures 4a and b), but no spacers targeted the intergenic regions of the two phage genomes. The majority of spacers with targets (49 and 39, respectively) were from the CRISPR-CasY systems in QZM Roizmanbacteria genomes, and all the other targeting spacers were from the Type I-B and III-A systems of C2-Gp7. No spacer from QZM_B4_Woesebacteria_36_36 (78 unique spacers) and the other 10 type III-A systems targeted the two complete phage genomes.

For phylogenetic analyses, we searched the NCBI database for capsid proteins similar to those in the genomes reported here (Figure 3) and identified a scaffold containing a similar capsid ORF that was binned into a Roizmanbacteria genome (Candidatus Roizmanbacteria bacterium RIFOXYA1_FULL_41_12; (Anantharaman et al. 2016)) (Supplementary Figure 2). Comparative analyses showed a close relationship between the sequences of this prophage and the two complete phages mentioned above, including homologies for the capsid and two terminase proteins. In addition, these genes and several other hypothetical proteins share gene arrangements (Supplementary Figure 6). Thus, we conclude the two phage genomes reported are the full sequences for lysogenic (temperate) phage found in Roizmanbacteria genomes.

### An unusual CRISPR-CasY system with a fragmented CasY effector and self-targeting spacers

Among the candidate CasY sequences from the Tibet hot springs predicted protein dataset were three adjacent partial proteins on a scaffold from sample GD2_1. In combination, the three open reading frames appear to comprise a fragmented CasY protein (defined as “fCasY“). We identified Cas1 and a CRISPR locus adjacent to the fCasY (Figure 5a). Read mapping to the scaffold revealed that the CasY was fragmented by two mutations. One involves deletion of A (from “AAAAA” to “AAAA“) and introduces a TAA stop codon five amino acids downstream. This mutation occurred in all the mapped reads, indicating that all the cells have CasY fragmented at this position. The second mutation is a single nucleotide substitution from “C” to “T”, which introduces a TAA stop codon. This mutation was detected in 82% of the mapped reads. Interestingly, however, the three conserved motifs (RuvC-I, -II and -III) are preserved in the largest protein fragment and all the catalytic residues are shared with functional CasY proteins (Figure 1b and Figure 5a). We identified the ribosome binding site (RBS) for fragments 1 and 2 as TAA, the same RBS associated with 353 of 946 ORFs of this Roizmanbacteria genome. The longest fragment is predicted to have a RBS of AAT, which was only shared by 55 ORFs.

The fCasY locus includes 22 unique spacers, six of which were detected only once in the mapped reads (Figure 5b). We reconstructed the CRISPR locus (Figure 5b) and found that all of the single copy spacers are at the locus end that is closest to the Cas1 protein. As in prior studies, we infer that these were recently added to the diversifying end of the CRISPR locus in a subset of cells. Interestingly, 12 out of the 22 unique spacers target the scaffolds of the C2-Gp5 genome, which encodes the fCasY system (Figure 5c, Supplementary Table 6). In detail, 11 spacers targeted Roizmanbacteria genes, including those encoding a PINc domain ribonuclease, two permeases, a sigma-70 RNA polymerase and three hypothetical proteins with transmembrane domains. Only one spacer matched an intergenic region, which is next to two tRNAs (His and Thr). This spacer was recently acquired, as it is encoded on three reads that also sampled part of the leader sequence (Figure 5b, Supplementary Table 6). Several of the self-targeting spacers are located in the old end of the locus (Figure 5b) and occurred in majority of the cells in the population. Thus, we infer that Roizmanbacteria with these self-targeting spacers have survived for a substantial period of time.

In addition to the fCasY locus, we identified type III-A and I-B CRISPR-Cas systems in the C2-Gp5 genome. Notably, one spacer from the type III-A and I-B systems and two fCasY spacers target a complete 34,706 bp phage genome GD2_3_Phage_34_19 (Supplementary Figure 7, Supplementary Table 6) assigned to *Podoviridae*. A Cas4-like protein was detected in this phage genome (Supplementary Figure 7). As phage with Cas4-like proteins can induce their hosts to acquire self-targeting spacers (Hooton and Connerton 2014), the presence of this protein may explain acquisition of self-targeting spacers by the C2-Gp5 genome.

Spacers from the loci of C2-Gp5 target other putative phage scaffolds (Supplementary Table 5). For example, one fCasY spacer targets GD2_3_scaffold_2486, which encodes a putative phage gene. Spacers from both fCasY and I-B systems target GD2_2_scaffold_18083, which encodes a phage tail tape measure protein. Two spacers from the type I-B system target GD2_3_scaffold_517, which encodes a capsid protein that is distantly related to that in the prophage of C2-Gp3 (Figure 3).

### PAMs 5‘-TA and 5’-TG are shared by systems with both CasY and fCasY

The PAM is used for the acquisition of spacers into the CRISPR array and is important for target recognition and cleavage (Hille et al. 2018). We determined the probable PAM of the CasY systems reported here to target the two complete phage genomes (QZM_A2_Phage_33_19 and QZM_B3_Phage_33_79). Among all the 39 unique target locations on these two phage genomes (88 spacers in total), 20 had a potential 5‘ TA PAM and 14 had a potential 5’ TG PAM (Supplementary Figure 8, Supplementary Table 6). Moreover, the one spacer in the CRISPR-CasY system of the C1-Gp1 genome that targets GD2_3_Phage_34_19 also has a 5’ TA PAM (Supplementary Figure 7). Previously, the PAM determined for the CasY.1 of *Candidatus* Katanobacteria using an *in vitro* approach was a 5‘ TA, and both 5’ TA (dominant) and 5’ TG PAMs occur, based on *in vivo* data (Burstein et al. 2017). For the fCasY, we checked to see if the self-targeting spacers have the same PAM as that of other CasY proteins. If this was not the case, the genomic region matching the spacer may not be recognized as a target by the fCasY CRISPR system. Among the 12 self-targeting spacers, 7 have 5‘ TA and 4 have 5’ TG PAMs and one has a possible 5’ AT PAM (Supplementary Table 6). Among the 5 fCasY spacers targets on phage scaffolds, two have 5‘ TA PAMs and two have 5’ TG PAMs.

In combination the results indicate that both general CasY proteins and fCasY in this study use the 5’ TA/TG PAM sequences for spacer acquisition and protospacer recognition. We identified a few targets with other PAM sequences (Supplementary Table 6), but it is possible that these targets have mutated the PAM sites during their evolutionary history, as previously documented (Paez-Espino et al. 2015).

### Potential phage-host genetic interactions

When examining the genomic context of CRISPR-CasY systems we noted four very short genes located next to the CRISPR array in the C2-Gp7 genome (Supplementary Figure 9). All four genes had at least one homologue in the three complete phage and one prophage (BLASTp e-value thresholds = 1e-5) and when two or more homologues were identified in the same genome, they were together. However, homologues were not identified in the other newly reconstructed and previously reported Roizmanbacteria genomes (Supplementary Table 4). The four genes in the C2-Gp7 genome and phage and prophage shared > 83% (up to 99%) nucleotide identity with > 80% alignment coverage, but none had a NCBI blast hit with similarity > 38% (> 50 alignment coverage). Given this, and the deduction that QZM_A2_Phage_33_19 and QZM_B3_Phage_33_79 infect C2-Gp7 Roizmanbacteria (based on CRISPR spacer targeting), we conclude that there may have been lateral transfer of novel proteins related to phage-host interactions between Roizmanbacteria and their phage.

## Discussion

CPR bacteria account for a huge amount of diversity within the Bacterial domain, but the mechanisms of their interactions with phage and the phage that infect them have remained largely undocumented. In part, this is due to scant information about their CRISPR-Cas systems, despite extensive genomic sampling from a wide variety of sites in nature (Burstein et al. 2016, 2017; Dudek et al. 2017; Castelle and Banfield 2018). In this study, we report an unexpected diversity of CRISPR-Cas systems in the genomes of bacteria from the CPR phylum of Roizmanbacteria, both from newly reconstructed sequences from multiple hot spring sediments of Tibet, China (Supplementary Table 1) and some previously published genomes. Most of them are CasY-based systems (Figure 1a, Table 1). These new sequences constrain more and less highly conserved regions of CasY proteins, information that may be important in future efforts directed at tailoring the properties of genome-editing enzymes.

The finding that some of the Roizmanbacteria genomes encode multiple CRISPR-Cas systems, including the relatively large types I-B and III-A, is unexpected, given the overall paucity of systems in CPR bacteria, and their small genome sizes (Figure 1b). We infer that these systems are mostly active, given the identification of targets on potential phage scaffolds and evidence for locus diversification. Considering that majority of the spacers with targets on the three complete phage and one prophage were from CRISPR-CasY systems (Figures 2d, and 4a and b), it seems that CasY is the primary CRISPR-Cas system used by these bacteria for phage defense. In the case of the Roizmanbacteria with only a degenerate Type III-A CRISPR-Cas system, defense may rely upon a restriction-modification system, as suggested previously for CPR bacteria that lack any CRISPR-Cas system (Burstein et al. 2016) (Figure 2). In support of this correlation, restriction-modification systems were not detected in those Roizmanbacteria with seemingly functional CRISPR-Cas systems (Figure 1a). The discovery of two copies Cas12a proteins in a single system of two genomes is an additional case of unexpected investment in CRISPR-Cas-based phage defense by CPR bacteria (Figure 1a, Supplementary Figure 2). Overall, the genomes of Roizmanbacteria contained three of the six types of CRISPR-Cas systems reported so far (i.e. type I, III and V), expanding our understanding of the investment of CPR bacteria in CRISPR-Cas-based defense.

The availability of a pool of CRISPR spacers enabled discovery of three Roizmanbacteria-infecting phage for which complete genomes were reconstructed, and one prophage (Figures 2-4, Supplementary Figure 7). These are the first reported phage infecting members of the Microgenomates superphylum of the CPR. All of these phage, along with the previously reported CPR phage, were assigned to *Podoviridae* and *Siphoviridae* of the *Caudovirales* order (Figure 3). The phylogenetic relatedness and genetic similarity among the *Podoviridae* phages obtained in this study and a Roizmanbacteria prophage deposited at NCBI (Figure 3, Supplementary Figure 6), and also the potential phage-host genetic interactions (Supplementary Figure 9), may indicate stable and similar host-phage relationships in a variety of habitats.

An interesting aspect of the CRISPR-CasY analyses was the fCasY system in one Roizmanbacteria that includes a locus with self-targeting spacers. It may be significant that a Cas4-like protein is encoded in the genome of a phage that replicates in this Roizmanbacteria, given that a Cas4-like protein in a *Campylobacter* sp. phage was suggested to facilitate acquisition of self-targeting spacers into the CRISPR-Cas system of its host (Hooton and Connerton 2014). Roizmanbacteria lack the RecBCD mediated double-stranded DNA break repair complex, the only documented mechanism for avoidance of self-targeting spacer acquisition (Levy et al. 2015). Thus, it is plausible that the phage-encoded Cas4-like protein led to acquisition of the self-targeting spacers, which should result in autoimmunity (Stern et al. 2010).

Autoimmunity can be avoided via loss of cas genes, mutated repeats adjacent to self-targeting spacers, extended base-pairing with the upstream flanking repeat, and the absence of a PAM in the chromosomal region matched by the spacer (Stern et al. 2010), none of which were observed here. Autoimmunity also could be countered via loss of cas gene function. Interestingly, the fCasY harboured conserved RuvC domains and catalytic residues found in intact CasY proteins (Figure 1b, Figure 5). However, given the relatively high abundance of Roizmanbacteria with fCasY in the community (1.37%), we infer that the fCasY protein fragmentation led to loss of cleavage function, preventing autoimmunity. It is possible that the region of the fCasY protein responsible for binding to the target sequence is encoded on a different gene fragment than that encoding the nuclease domain, so that the CRISPR RNA does not recruit the protein fragment with nuclease function.

The presence of old end CRISPR locus spacers that target the host chromosome suggests that the fCasY has been present in the genomes of the Roizmanbacteria C2-Gp5 population for some time. Why has this gene, or the entire locus, not been lost? It is possible that the spacers of the fCasY locus retain some function, for example in gene regulation (possibly involving binding of CRISPR RNAs to the DNA during transcription). Experiments will be required to determine whether fragments of fCasY can reassemble and bind to the genomic regions targeted by the self-targeting spacers (without cleavage) and to determine if the spacer-directed binding domain is on fragment 1 or 2 (Figure 5a).

In conclusion, CRISPR-Cas systems are unexpectedly common in a subset of CPR bacteria, and the number, variety and potential functional diversity of these systems is greater than expected. It is already established that CRISPR-CasY systems from these intriguing and enigmatic bacteria will have biotechnological value. Lessons from natural system studies such as reported here may provide information about CasY sequence variety and function that may be useful in enzyme engineering. Beyond this, the new information about CPR bacteria, their phage and the mechanisms of their interactions expands our understanding of the complex phenomena that shape the structure and functioning of natural microbial communities.

## Author contributions

L.X.C. and J.F.B. designed the study. W.J.L. supported for the metagenomic sequencing. L.X.C. performed the metagenomic assembly, HMM search and scaffold extension and curation. L.X.C. and J.F.B. performed genome binning and curation. L.X.C. and J.F.B. conducted data analyses with input from B.A.S. and R.M.. L.X.C. and J.F.B. wrote the manuscript. All authors read and approved the final manuscript.

## Acknowledgement

This research was supported by the Microbiology Program of the Innovative Genomics Institute. W.J.L. was financially supported by the Science and Technology Infrastructure work project (No. 2015FY110100), and the Natural Science Foundation of Guangdong Province, China (No. 2016A030312003). B.A.S is supported by the National Science Foundation Graduate Research Fellowship (DGE 1752814).

## Conflict of interest

J.F.B. is a founder of Metagenomi. J.A.D. is a co-founder of Caribou Biosciences, Editas Medicine, Intellia Therapeutics, Scribe Therapeutics, and Mammoth Biosciences. J.A.D. is a scientific advisory board member of Caribous Biosciences, Intellia Therapeutics, eFFECTOR Therapeutics, Scribe Therapeutics, Synthego, Metagenomi, Mammoth Biosciences and Inari. J.A.D is a member of the board of directors at Driver and Johnson & Johnson and has sponsored research projects by Roche Biopharma and Biogen.

## References

Anantharaman, Karthik, Christopher T. Brown, Laura A. Hug, Itai Sharon, Cindy J. Castelle, Alexander J. Probst, Brian C. Thomas, et al. 2016. “Thousands of Microbial Genomes Shed Light on Interconnected Biogeochemical Processes in an Aquifer System.” Nature Communications 7:13219.

Andersson Anders F., and Jillian F. Banfield. 2008. “Virus Population Dynamics and Acquired Virus Resistance in Natural Microbial Communities.” Science 320 (5879): 1047–50.

Bankevich, Anton, Sergey Nurk, Dmitry Antipov, Alexey A. Gurevich, Mikhail Dvorkin, Alexander S. Kulikov, Valery M. Lesin, et al. 2012. “SPAdes: A New Genome Assembly Algorithm and Its Applications to Single-Cell Sequencing.” Journal of Computational Biology: A Journal of Computational Molecular Cell Biology 19 (5): 455–77.

Brown Christopher T., Laura A. Hug, Brian C. Thomas, Itai Sharon, Cindy J. Castelle, Andrea Singh, Michael J. Wilkins, Kelly C. Wrighton, Kenneth H. Williams, and Jillian F. Banfield. 2015. “Unusual Biology across a Group Comprising More than 15% of Domain Bacteria.” Nature 523 (7559): 208–11.

Burstein, David, Lucas B. Harrington, Steven C. Strutt, Alexander J. Probst, Karthik Anantharaman, Brian C. Thomas, Jennifer A. Doudna, and Jillian F. Banfield. 2017. “New CRISPR-Cas Systems from Uncultivated Microbes.” Nature 542 (7640): 237–41.

Burstein, David, Christine L. Sun, Christopher T. Brown, Itai Sharon, Karthik Anantharaman, Alexander J. Probst, Brian C. Thomas, and Jillian F. Banfield. 2016. “Major Bacterial Lineages Are Essentially Devoid of CRISPR-Cas Viral Defence Systems.” Nature Communications 7 (February): 10613.

Capella-Gutiérrez, Salvador, José M. Silla-Martínez, and Toni Gabaldón. 2009. “trimAl: A Tool for Automated Alignment Trimming in Large-Scale Phylogenetic Analyses.” Bioinformatics 25 (15): 1972–73.

Castelle Cindy J., and Jillian F. Banfield. 2018. “Major New Microbial Groups Expand Diversity and Alter Our Understanding of the Tree of Life.” Cell 172 (6): 1181–97.

Castelle Cindy J., Christopher T. Brown, Karthik Anantharaman, Alexander J. Probst, Raven H. Huang, and Jillian F. Banfield. 2018. “Biosynthetic Capacity, Metabolic Variety and Unusual Biology in the CPR and DPANN Radiations.” Nature Reviews. Microbiology 16 (10): 629–45.

Chen Janice S., and Jennifer A. Doudna. 2017. “The Chemistry of Cas9 and Its CRISPR Colleagues.” Nature Reviews Chemistry 1 (10): 0078.

Crooks G. E., G. Hon, J. M. Chandonia, and S. E. Brenner. 2004. “WebLogo: A Sequence Logo Generator.” Genome Research 14 (6): 1188–90.

Dick Gregory J., Anders F. Andersson, Brett J. Baker, Sheri L. Simmons, Brian C. Thomas, A. Pepper Yelton, and Jillian F. Banfield. 2009. “Community-Wide Analysis of Microbial Genome Sequence Signatures.” Genome Biology 10 (8): R85.

Dudek Natasha K., Christine L. Sun, David Burstein, Rose S. Kantor, Daniela S. Aliaga Goltsman, Elisabeth M. Bik, Brian C. Thomas, Jillian F. Banfield, and David A. Relman. 2017. “Novel Microbial Diversity and Functional Potential in the Marine Mammal Oral Microbiome.” Current Biology: CB 27 (24): 3752–62.e6.

Eddy S. R. 1998. “Profile Hidden Markov Models.” Bioinformatics 14 (9): 755–63.

Edgar, Robert C. 2004. “MUSCLE: Multiple Sequence Alignment with High Accuracy and High Throughput.” Nucleic Acids Research 32 (5): 1792–97.

Grissa, Ibtissem, Patrick Bouchon, Christine Pourcel, and Gilles Vergnaud. 2008. “On-Line Resources for Bacterial Micro-Evolution Studies Using MLVA or CRISPR Typing.” Biochimie 90 (4): 660–68.

Hille, Frank, Hagen Richter, Shi Pey Wong, Majda Bratovič, Sarah Ressel, and Emmanuelle Charpentier. 2018. “The Biology of CRISPR-Cas: Backward and Forward.” Cell 172 (6): 1239–59.

Hooton, Steven P. T., and Ian F. Connerton. 2014. “Campylobacter Jejuni Acquire New Host-Derived CRISPR Spacers When in Association with Bacteriophages Harboring a CRISPR-like Cas4 Protein.” Frontiers in Microbiology 5:744.

Hug Laura A., Brett J. Baker, Karthik Anantharaman, Christopher T. Brown, Alexander J. Probst, Cindy J. Castelle, Cristina N. Butterfield, et al. 2016. “A New View of the Tree of Life.” Nature Microbiology 1 (April): 16048.

Hyatt, Doug, Gwo-Liang Chen, Philip F. Locascio, Miriam L. Land, Frank W. Larimer, and Loren J. Hauser. 2010. “Prodigal: Prokaryotic Gene Recognition and Translation Initiation Site Identification.” BMC Bioinformatics 11 (March): 119.

Langmead, Ben, and Steven L. Salzberg. 2012. “Fast Gapped-Read Alignment with Bowtie 2.” Nature Methods 9 (4): 357–59.

Letunic I., and P. Bor. 2006. “Interactive Tree Of Life (iTOL): An Online Tool for Phylogenetic Tree Display and Annotation.” Bioinformatics 23 (1): 127–28.

Levy, Asaf, Moran G. Goren, Ido Yosef, Oren Auster, Miriam Manor, Gil Amitai, Rotem Edgar, Udi Qimron, and Rotem Sorek. 2015. “CRISPR Adaptation Biases Explain Preference for Acquisition of Foreign DNA.” Nature 520 (7548): 505–10.

Luef, Birgit, Kyle R. Frischkorn, Kelly C. Wrighton, Hoi-Ying N. Holman, Giovanni Birarda, Brian C. Thomas, Andrea Singh, et al. 2015. “Diverse Uncultivated Ultra-Small Bacterial Cells in Groundwater.” Nature Communications 6 (1). https://doi.org/10.1038/ncomms7372.

Nuñez James K., Philip J. Kranzusch, Jonas Noeske, Addison V. Wright, Christopher W. Davies, and Jennifer A. Doudna. 2014. “Cas1–Cas2 Complex Formation Mediates Spacer Acquisition during CRISPR–Cas Adaptive Immunity.” Nature Structural & Molecular Biology 21 (6): 528–34.

Olm Matthew R., Christopher T. Brown, Brandon Brooks, and Jillian F. Banfield. 2017. “dRep: A Tool for Fast and Accurate Genomic Comparisons That Enables Improved Genome Recovery from Metagenomes through de-Replication.” The ISME Journal 11 (12): 2864–68.

Paez-Espino, David, Emiley A. Eloe-Fadrosh, Georgios A. Pavlopoulos, Alex D. Thomas, Marcel Huntemann, Natalia Mikhailova, Edward Rubin, Natalia N. Ivanova, and Nikos C. Kyrpides. 2016. “Uncovering Earth’s Virome.” Nature 536 (7617): 425–30.

Paez-Espino, David, Itai Sharon, Wesley Morovic, Buffy Stahl, Brian C. Thomas, Rodolphe Barrangou, and Jillian F. Banfield. 2015. “CRISPR Immunity Drives Rapid Phage Genome Evolution in Streptococcus Thermophilus.” mBio 6 (2). https://doi.org/10.1128/mBio.00262-15.

Parks Donovan H., Christian Rinke, Maria Chuvochina, Pierre-Alain Chaumeil, Ben J. Woodcroft, Paul N. Evans, Philip Hugenholtz, and Gene W. Tyson. 2017. “Recovery of Nearly 8,000 Metagenome-Assembled Genomes Substantially Expands the Tree of Life.” Nature Microbiology 2 (11): 1533–42.

Pruesse, Elmar, Jörg Peplies, and Frank Oliver Glöckner. 2012. “SINA: Accurate High-Throughput Multiple Sequence Alignment of Ribosomal RNA Genes.” Bioinformatics 28 (14): 1823–29.

Pruesse, Elmar, Christian Quast, Katrin Knittel, Bernhard M. Fuchs, Wolfgang Ludwig, Jörg Peplies, and Frank Oliver Glöckner. 2007. “SILVA: A Comprehensive Online Resource for Quality Checked and Aligned Ribosomal RNA Sequence Data Compatible with ARB.” Nucleic Acids Research 35 (21): 7188–96.

Schulz, Frederik, Emiley A. Eloe-Fadrosh, Robert M. Bowers, Jessica Jarett, Torben Nielsen, Natalia N. Ivanova, Nikos C. Kyrpides, and Tanja Woyke. 2017. “Towards a Balanced View of the Bacterial Tree of Life.” Microbiome 5 (1): 140.

Shmakov, Sergey, Omar O. Abudayyeh, Kira S. Makarova, Yuri I. Wolf, Jonathan S. Gootenberg, Ekaterina Semenova, Leonid Minakhin, et al. 2015. “Discovery and Functional Characterization of Diverse Class 2 CRISPR-Cas Systems.” Molecular Cell 60 (3): 385–97.

Song, Zhao-Qi, Feng-Ping Wang, Xiao-Yang Zhi, Jin-Quan Chen, En-Min Zhou, Feng Liang, Xiang Xiao, et al. 2012. “Bacterial and Archaeal Diversities in Yunnan and Tibetan Hot Springs, China.” Environmental Microbiology 15 (4): 1160–75.

Stamatakis Alexandros. 2014. “RAxML Version 8: A Tool for Phylogenetic Analysis and Post-Analysis of Large Phylogenies.” Bioinformatics 30 (9): 1312–13.

Stern, Adi, Leeat Keren, Omri Wurtzel, Gil Amitai, and Rotem Sorek. 2010. “Self-Targeting by CRISPR: Gene Regulation or Autoimmunity?” Trends in Genetics: TIG 26 (8): 335–40.

Westra Edze R., Stineke van Houte, Sam Oyesiku-Blakemore, Ben Makin, Jenny M. Broniewski, Alex Best, Joseph Bondy-Denomy, Alan Davidson, Mike Boots, and Angus Buckling. 2015. “Parasite Exposure Drives Selective Evolution of Constitutive versus Inducible Defense.” Current Biology: CB 25 (8): 1043–49.

Yarza, Pablo, Pelin Yilmaz, Elmar Pruesse, Frank Oliver Glöckner, Wolfgang Ludwig, Karl-Heinz Schleifer, William B. Whitman, Jean Euzéby, Rudolf Amann, and Ramon Rosselló-Móra. 2014. “Uniting the Classification of Cultured and Uncultured Bacteria and Archaea Using 16S rRNA Gene Sequences.” Nature Reviews. Microbiology 12 (9): 635–45.

Zetsche, Bernd, Jonathan S. Gootenberg, Omar O. Abudayyeh, Ian M. Slaymaker, Kira S. Makarova, Patrick Essletzbichler, Sara E. Volz, et al. 2015. “Cpf1 Is a Single RNA-Guided Endonuclease of a Class 2 CRISPR-Cas System.” Cell 163 (3): 759–71.

